# Haptic metameric textures

**DOI:** 10.1101/653550

**Authors:** Scinob Kuroki, Masataka Sawayama, Shin’ya Nishida

## Abstract

Humans sense spatial patterns through their eyes and hands. Past studies have revealed differences (as well as similarities) between vison and touch in texture processing (e.g., eye is good at detecting texture boundaries, while hand can discriminate subtle texture differences), but the underlying computational differences remains poorly understood. Here we transcribed various textures as surface relief patterns by 3D-printing, and analyzed the tactile discrimination performance regarding the sensitivity to image statistics. The results suggest that touch is sensitive to texture differences in lower-order statistics (e.g., statistics of local amplitude spectrum), while may not to those in the higher-order statistics (e.g., joint statistics of local orientations). In agreement with this, we found that pairs of synthesized textures differing only in higher-order statistics were nearly indiscriminable (metameric) by touch, while easily discriminable by vision. Our findings show that touch and vision sense spatial information using different and complementary computational strategies.

## Introduction

Most objects in the world are covered by surfaces with a variety of textures. By sensing surface textures, we, and many other animals, are able to specify distinct surface areas, identify materials, and estimate surface conditions. Touch and vision are the two main sensory modalities contributing to surface texture perception, and several studies have highlighted similarities and differences in their abilities to access various spatial properties (see review for Klatzky et al., 2013, Bensmaia 2009, Whitaker et al., 2008). For example, vision is good at detecting geometric patterns such as texture boundaries (Klatzky et al., 2013, Lederman & Abbott 1981), while touch can detect invisibly small surface texture properties (Bergmann Tiest and Kappers, 2007; Heller, 1989; LaMotte & Srinivasan, 1991). These differences between the two modalities can be partially ascribed to the differences in sensing hardware characteristics (e.g., higher sampling density for vision, availability of temporal vibration information for touch), but whether and how they differ each other in the higher neural spatial processing remains obscure. Here we try to show the difference in computation between the tactile and visual texture processing with regard to the sensitivity to image statistics.

Early studies on visual texture segregation revealed that the condition for two adjacent textures to be perceptually segregated is the presence of significant differences in the histogram of local orientation and spatial frequency (Gabor wavelets) (Julesz, 1962; Bergen & Adelson, 1988), which is presumably represented by the response distribution of V1 neurons (level #2 in Fig. 1). Recent studies examining the perceptual discrimination of natural textures further suggest that two textures are perceptually indistinguishable (become a metameric pair) in peripheral vision when they are matched not only in terms of V1 image statistics, but also in terms of the joint statistics of V1 responses (level #3 in Fig. 1), to which V2 and the higher cortical areas are responsive (Freeman & Simoncelli, 2011; Freeman et al., 2013; Ziemba et al., 2016). Human observers can visually discriminate V2 metamer textures (with identical Gabor and joint statistics), but only when using elaborated attentive spatial processing by central vision (Freeman et al., 2013; Rosenholtz, 2016).

**Fig. 1.**
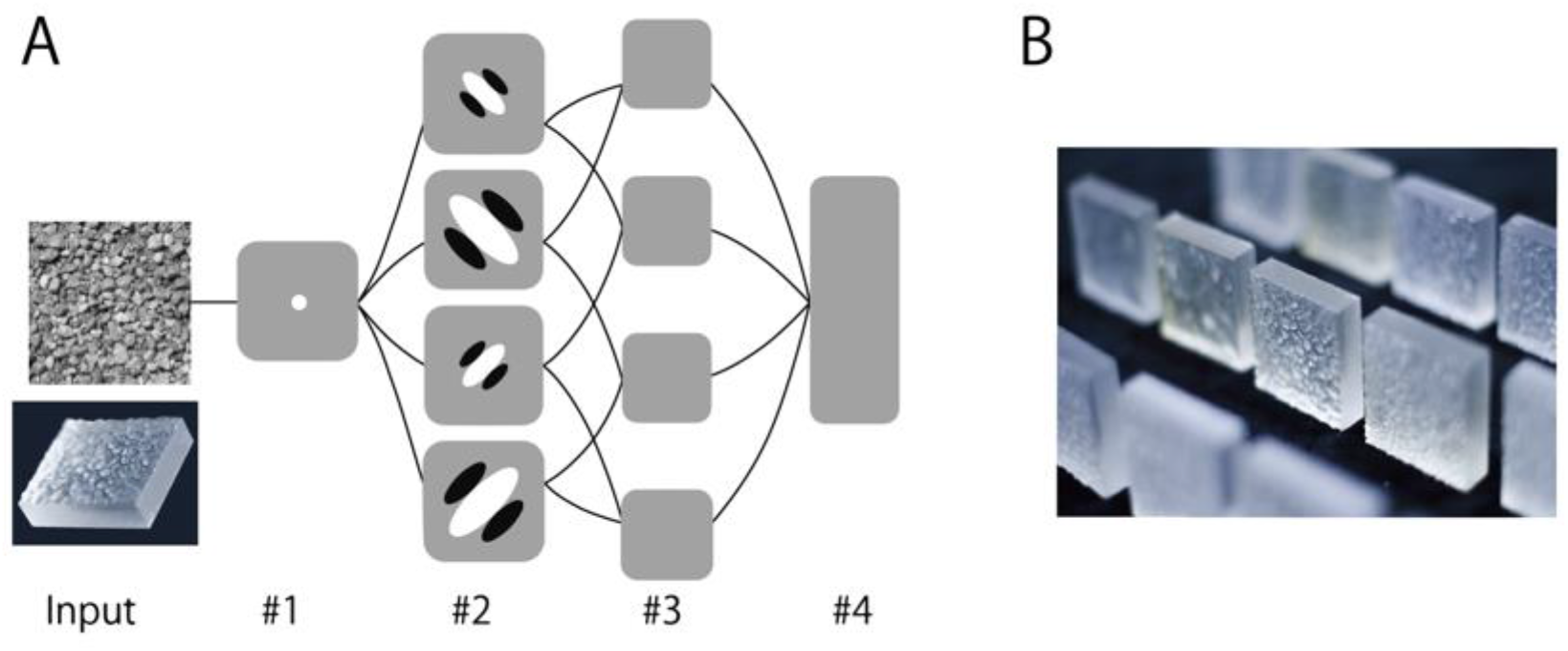
(A) Hypothetical diagram of hierarchical information processing in touch, inspired by visual processing. (B) 3D printed stimuli used for experiments.

In this paper, we will call the image statistics represented at or below the level #2 in Fig. 1 as lower order, and those represented at or beyond the level #3 as higher order. The lower-order statistics include the intensity histogram and Gabor statistics, and are associated with the Fourie amplitude spectrum and the second-order statistics in the terminology of Julesz (1962), while the higher-order statistics include joint statistics (Freeman & Simoncelli, 2011), and are associated with the Fourier phase spectrum. According to this terminology, visual texture perception is sensitive to higher-order statistics, in addition to lower-order statistics. The question is whether this is also the case for tactile texture perception.

The somatosensory system has a spatial-information processing stream analogous to that of the visual system. First, a spatiotemporal pattern of skin deformation is sampled by mechanoreceptors. The signals from the mechanoreceptors are then pooled, suppressed by surrounds, and sent to the cortex via peripheral afferent neurons. Some peripheral afferents may be able to carry some orientation information (Delhaye et al., 2018; Pruszynski & Johansson, 2014), but orientation selectivity is much more common and robust in the primary somatosensory cortex (S1, area 3b) (e.g., Bensmaia et al., 2008). The tactile receptive field of S1 neurons can be approximated by Gabor functions (DiCarlo et al., 1998, 2000), as are the visual receptive field of V1 neurons. Somatosensory processing becomes more elaborate beyond S1 (e.g., Thakur et al., 2006). For example, some neurons in S2 have selectivity for higher-order shape features, such as the curvature of line stimuli, similar to that of visual neurons in V4 (Yau et al., 2009).

Here we investigated the performance of the tactile system in discriminating a variety of texture patterns (artificial textures made of numerous Gabors, and natural textures). All the patterns were easy to discriminate by vision. Through the analysis of the tactile discrimination performance, we considered what sorts of texture differences the tactile system is sensitive to, and, more specifically, whether it can utilize higher-order image statistics (at or beyond level #3 in Fig. 1) as does visual texture perception. There is little psychophysical evidence that tactile texture discrimination is sensitive to higher-order image statistics. The major spatial property of tactile texture perception intensively investigated in the past is roughness (Bensmaia, 2009; Klatzky et al., 2013; Hollins et al., 1993, 2000; Tiest, 2010; Tiest and Kapers, 2006). (Other properties, such as hardness and stickiness, are not purely spatial.) The stimulus spatial parameters that are known to affect roughness, such as the spatial period and inter-ridge spacing (e.g. Goodwin et al., 1989; Lederman, 1983; Lederman et al., 1972; Sathian et al., 1989, Taylor and Lederman, 1975), can be described in terms of lower-order image statistics. However, the neural responses to shapes/curvatures (Yau et al., 2009, 2013) suggest potential sensitivity to higher-order statistics.

Most previous tactile texture studies used relatively simple artificial stimuli, such as dots, gratings (Bensmaia and Hollins, 2005; Goodwin et al., 1989; Hollins and Bensmaia, 2007; Lederman, 1983; Lederman and Taylor, 1972; Sathian et al., 1989, Taylor and Lederman, 1975), or went with daily natural surfaces, such as fabric, wood, metal (Weber et al., 2013; Yokosaka et al., 2017). With these stimuli, it is not easy to examine the contribution of higher-order image statistics separately from those of lower-order ones. Here we overcame this limitation using a high-resolution 3D printer. By carving the surface of a sample material with the printer, we transcribed complex patterns including natural image textures into patterns of surface depth modulation. By manipulating the image statistics of the printed patterns, we were able examined whether tactile texture perception can utilize higher-order texture statistics.

## Results

We evaluated human tactile accuracy in discriminating a pair of 3D-printed textures using an ABX task (Kingdom & Prins, 2010). After passively scanning three textures, A, B, and X in order, with the fingertip of the right index finger, the observer had to judge whether X was a 180°-rotated version of A or B (Fig. 2B). Since X was rotated, observers could not perform the task based on simple pattern matching. The maximum depth of the surface texture was approximately 2 mm, which was well above the minimum detectable depth magnitude (Bolanowski et al., 1988; Gescheider et al., 2001, 2002). Most previous studies on tactile texture perception asked observers to judge a specific roughness feature (e.g., “rate the roughness from 0 to 9” or “report which one was rougher”). In contrast, our method allowed us to account for any perceptual feature, including roughness, that the observers could use to discriminate stimuli. Furthermore, it enabled us to find two physically different textures that are ‘metameric’ (perceptually indiscriminable in any way).

**Fig. 2.**
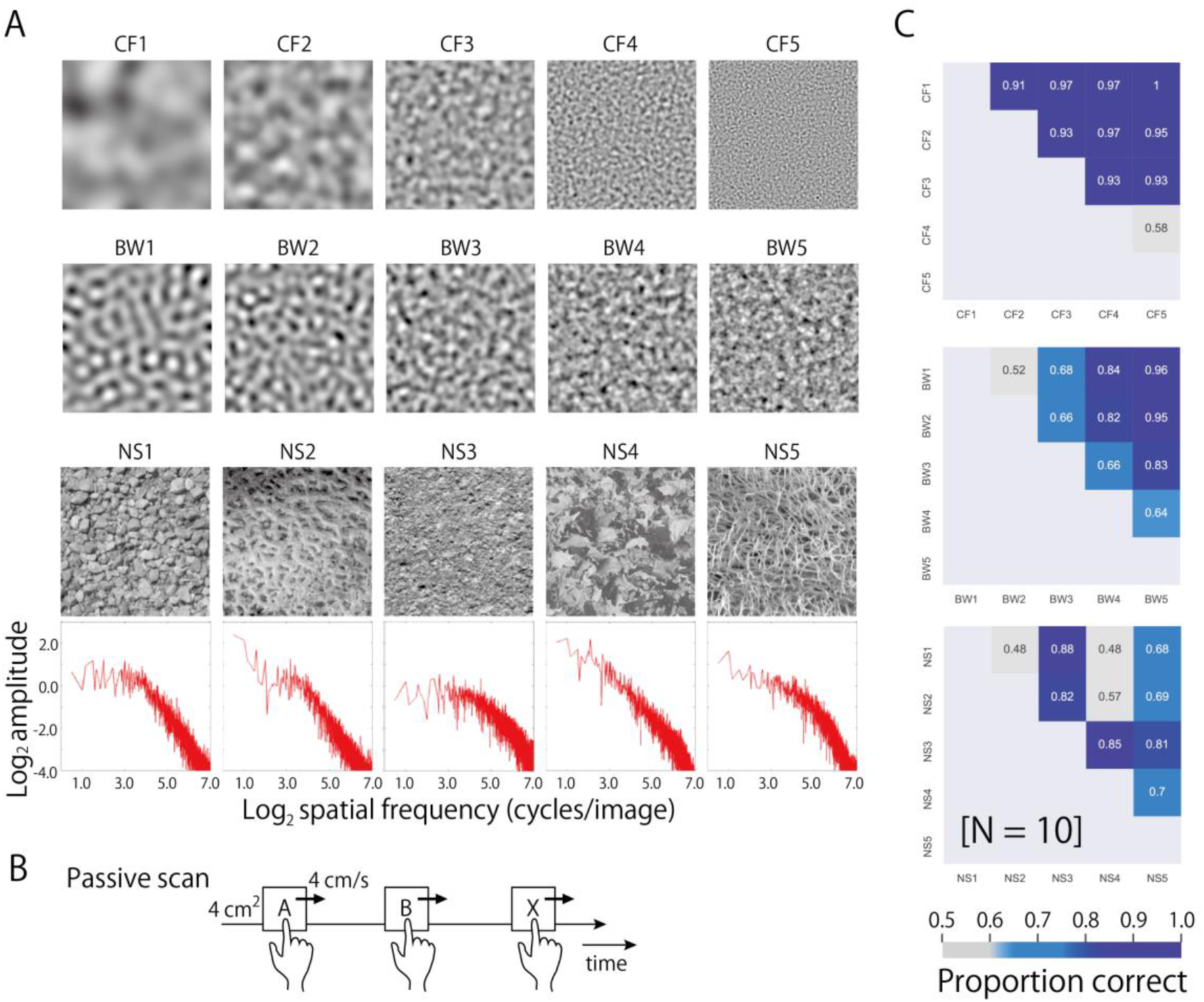
ABX texture discrimination task for 3D-printed stimuli. (A) Visual images (256×256 pixels) were converted into 3D carvings (40×40×10-12 mm) by taking intensity values as height maps. The intensity level (the mean and standard deviation) was matched across images. The bottom panels are log amplitude spectra of NS images (i.e., the amplitude of each spatial frequency component in the Fourier spectrum, averaged across orientations. See also Fig. 4A for details.) (B) Time course of the ABX experiment. Passive scan condition. Observers placed their index finger on the first resting cushion and the linear stage started to move at 40 mm/sec. After the finger contacted the tactile stimulus and swiped for one second, the stage stopped on the next resting cushion. This was repeated three times, once each for an A, B and X stimulus, with X a rotated version of A or B. Observers were asked to report verbally whether the third stimulus was the first or second one. No feedback signal was provided. (C) Results. Discrimination performance (proportion correct) for each stimulus pair is shown by numbers and colors in a matrix format. The 95% confidence interval of the chance performance is 0.42-0.57 (uncorrected), and 0.38-0.62 (Bonferroni corrected).

**Fig. 3.**
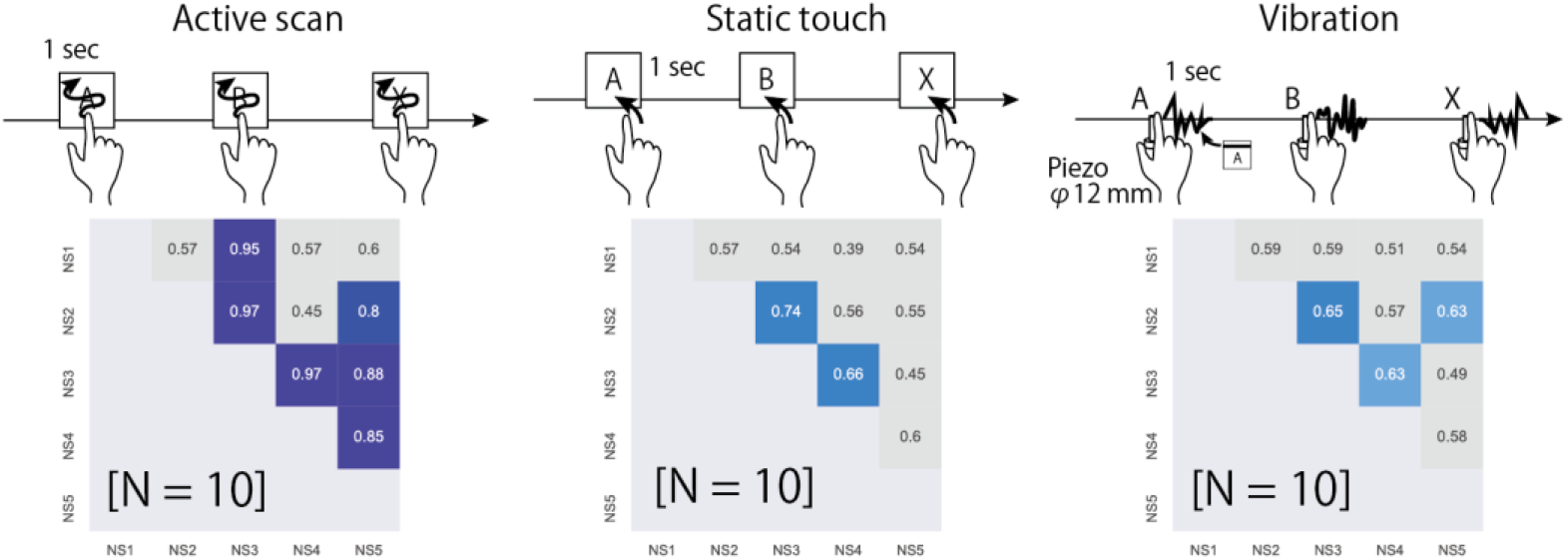
Texture discrimination for NS stimuli with the other touching modes. In the active scan condition, observers could freely explore the stimuli for one second with their index finger. In the static touch condition, observers put their finger on the stimuli for one second. They were not allowed to tangentially move/scan their finger over the stimuli. In the vibration condition, observers’ finger was vibrated by a piezo-electric actuator (MU120, MESS-TEK, Japan). The vibration pattern was one of the texture height profiles swept along a horizontal line. The observers could not discriminate some pairs of NS stimuli regardless of the touching mode. The 95% confidence interval of the chance performance is 0.42-0.57 (uncorrected), and 0.38-0.62 (Bonferroni corrected).

### Texture discrimination for 3D-printed stimuli

In the first series of experiments, discrimination performance was measured for three sets of five textures (Fig. 2A). The first two sets were spatially bandpass random noise patterns, each made of numerous Gabor components. The variables across textures were the center frequency (CF) for the first set and the bandwidth (BW) for the second set. The last set consisted of five natural visual textures (hereafter referred to as natural scenes (NS)).

The discrimination performance of ten observers obtained with the CF-variable set was very good (Fig. 2C, top panel): a one-octave difference in the CF was sufficient for nearly perfect discrimination, except for the high-frequency pair (CF4 and CF5). The results are consistent with previous findings obtained with analogous conditions (e.g., Bensmaia, 2009; Klatzky et al., 2013; Tiest, 2010). Varying the BW, on the other hand, is a novel approach in the quest for haptic sensitive stimulus variables. While discrimination accuracy of a one-octave difference in BW was 0.66 at best, that of a three-octave difference attained 0.95 (Fig. 2C, middle panel). Variations in the CF and BW are changes in lower-order image statistics (visible in the amplitude spectrum). The results therefore indicate that the tactile texture perception is sensitive to differences in low-order image statistics, although the tactile discrimination was not as good as the visual one (which is nearly perfect for the gray-level version of the stimuli).

For the third set, we printed monochromatic natural visual scene images (i.e., irregular patterns of stones, leaves, actiniae, etc. Fig. 2A), with transforming image intensity to carving depth. We used visual textures because we wanted to make direct comparison of the processing in the two modalities, and because we wanted a variety of textures with complex higher-order structures. Note also that our procedure did not produce highly unnatural tactile textures, since spatial coordinates are common between visual and tactile images, and the intensity-depth transformation has a theoretical ground such that deeper parts of surface texture are darker under diffuse illumination, known as vignetting (Koenderink, J., & van Doorn, A., 2003). We matched across textures the average and variance of the intensity (carving depth), while leaving the other statistical differences intact. Visually, five textures had very different spatial patterns. Nevertheless, the ten observers could barely discriminate most of the pairs by touch (Fig. 2C, bottom). Anecdotal reports from our observers suggest that the texture pattern (i.e., how the stimuli would look) was hard to guess and indistinguishable from other stimuli. The exception was NS3, which they described as somewhat ‘spikier’ than the rest. In summary, the same observers who could discriminate CF and BW stimuli could not discriminate most of the NS stimuli. Supplemental Fig. S1 shows the multi-dimensional scaling (MDS) of additional pairwise similarity judgments with CF, BW, and NS stimuli. In agreement with the results of the main discrimination experiment, NS stimuli are clustered in Fig. S1.

### Texture discrimination in different touching modes

Since haptic performance is known to be significantly affected by the touching mode, we repeated our ABX experiment for NS stimuli with three other touching modes. In the main experiment, we used the passive scan mode to match the speed and trajectory of scanning across different stimuli and different observers as much as possible. On the other hand, active scanning, in which observers can freely move their finger to explore the stimulus surface, may be able to provide richer spatial information than passive scanning (Kenshalo, 1978; Paillard et al., 1978). We therefore conducted the ABX experiment with the active scan mode. While the performance was slightly improved (average probability of 0.70 for passive scan; 0.76 for active scan, Fig. 3 left panel), the pattern of the results was similar to that in the passive scan condition. This is in good agreement with previous findings that tactile texture perception, including that of roughness and orientation, is relatively insensitive to changes in the exploration speed (Johnson and Yoshioka, 2002; Taylor and Lederman, 1975) and exploration method (Heller, 1989; Lamb, 1983; Olczak et al., 2018; Verrillo et al., 1999; Yoshioka et al., 2011). The other two modes were concerned with possible summation effects. Since the tactile system shows drastic spatial and temporal summation particularly with sub-threshold stimuli (Gescheider et al., 1999, 2005), texture discrimination performance might be seriously violated for complex textures like NS that give rapidly changing input in space and time. To reduce potentially negative effects of temporal/spatial summation on texture perception, we tested the static touch mode, where temporal information was limited, and the vibration mode, where spatial information was limited. In neither case did performance improve (Fig. 3, middle and right panels). In summary, the results indicate that some NS pairs are metameric regardless of the mode of touching. These trends across touching modes were the same for CF and BW stimuli (See Supplemental Fig. S2).

### Lower-order statistics and behavioural performance

The results obtained with the CF and BW stimuli indicate that tactile texture perception is sensitive to (some sorts of) differences in the amplitude spectrum or in the lower-order statistics. We next consider how well the discrimination performance for the NS stimuli can be explained by the differences in the amplitude spectrum. It is known that natural visual textures tend to have the amplitude spectrum falling with the spatial frequency by a factor of f^−a^ (Field, 1987). Our NS stimuli also have such amplitude spectra, which are similar to one another (Fig. 2A), although the slope of the spectrum differs for some textures. To analyze whether the similarity of amplitude spectra can explain the discrimination performance of tactile textures, we first integrated the amplitude differences between the paired textures over frequency (Fig. 4A) and then regressed the net amplitude difference to the discrimination performance by using a logistic regression analysis for all CF, BW, and NS conditions. Figure 4B shows the estimated performance plotted against the human performance. Although we did not consider orientation, the correlation was fairly high: R^2^=0.65 and 0.81 for the overall correlation and NS condition correlation, respectively. That is, the more similar the amplitude spectrum was, the more difficult the tactile texture discrimination became.

**Fig. 4.**
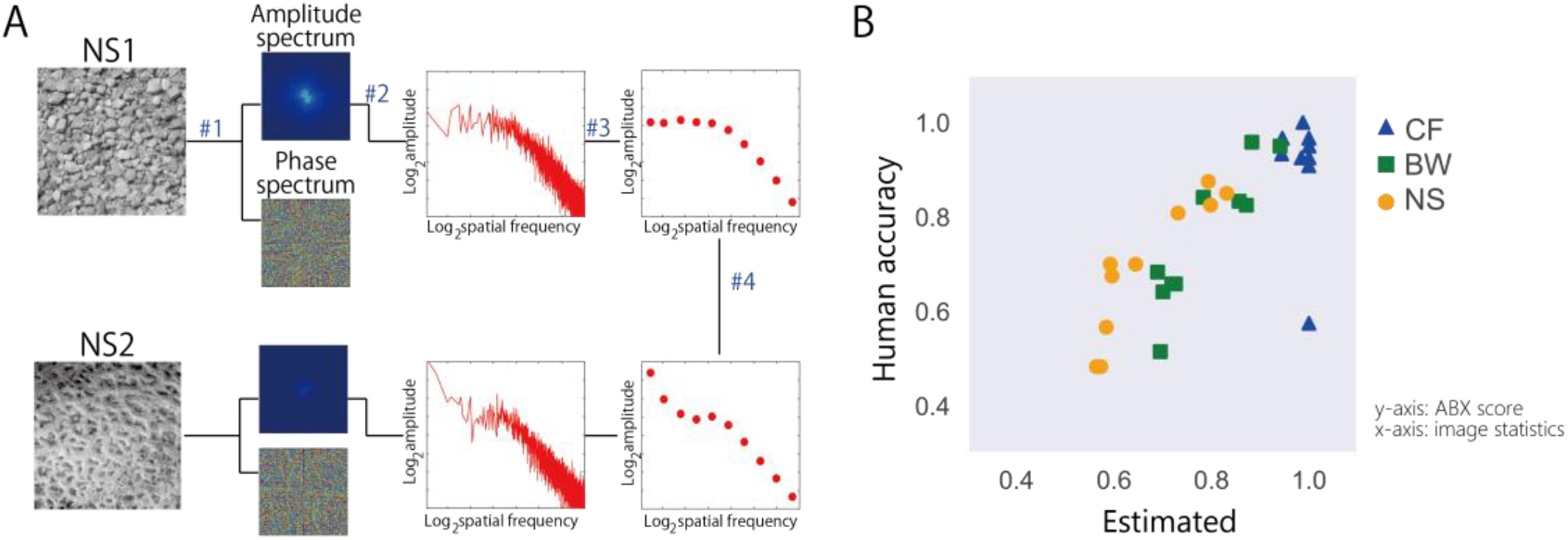
The discrimination performance for the natural visual texture can be explained by the amplitude difference. (A) Calculation of the amplitude difference. Step1 [#1]: The amplitude spectrum of each texture image is calculated through a 2D fast Fourier transformation (FFT). Step 2 [#2]: The amplitude of each spatial frequency component is averaged across different orientations on a log scale. Step 3 [#3]: The amplitude spectrum is sampled at constant intervals. Step 4 [#4]: The differences between the amplitude spectra of paired textures are used to explain the discrimination performance in Fig. 2. (B) Correlation between the human discrimination performance in the passive scan condition and the performance estimated by logistic regression analysis based on amplitude differences.

### Texture discrimination for stimuli with matched lower-order statistics

The results obtained so far can be ascribed solely to tactile texture processing sensitive to lower-order statistics, suggesting a hypothesis that tactile texture processing is unable to use higher-order statistics (e.g., joint Gabor statistics, phase spectra), to which visual texture perception is sensitive. This hypothesis predicts that textures with identical lower-order (subband) statistics are haptically indistinguishable (i.e., become a metameric pair) even if they differ from one another in the higher-order statistics. To test this, we made five images with identical subband histograms (Fig. 5A, bottom panels M1-M5) by matching the subband histogram of four NS images (NS2-NS5) to that of NS1 (based on Heeger and Bergen, 1995, see Methods). Visually, the matched images looked different from one another, and they were similar to the original images with regard to global patterning. However, haptic discrimination performance was nearly chance, 0.53 on average, and 0.61 at best. The result of the MDS analysis of the pairwise similarity ratings also indicate haptic metamerism of the matched stimuli: NS stimuli are similar but somehow distributed in perceptual space, while they are concentrated around the base stimulus (NS1) when their histograms are matched (See Supplemental Fig. S3). In agreement with the trends found with the group average, individual discrimination performance for the matched stimuli (M1-M5) was not significantly different from chance for all but one observers (9/10). This was in marked contrast to the original condition (NS1-NS5) where discrimination was significantly different from chance for all observers (10/10) (See Supplemental Table S1).

**Fig. 5.**
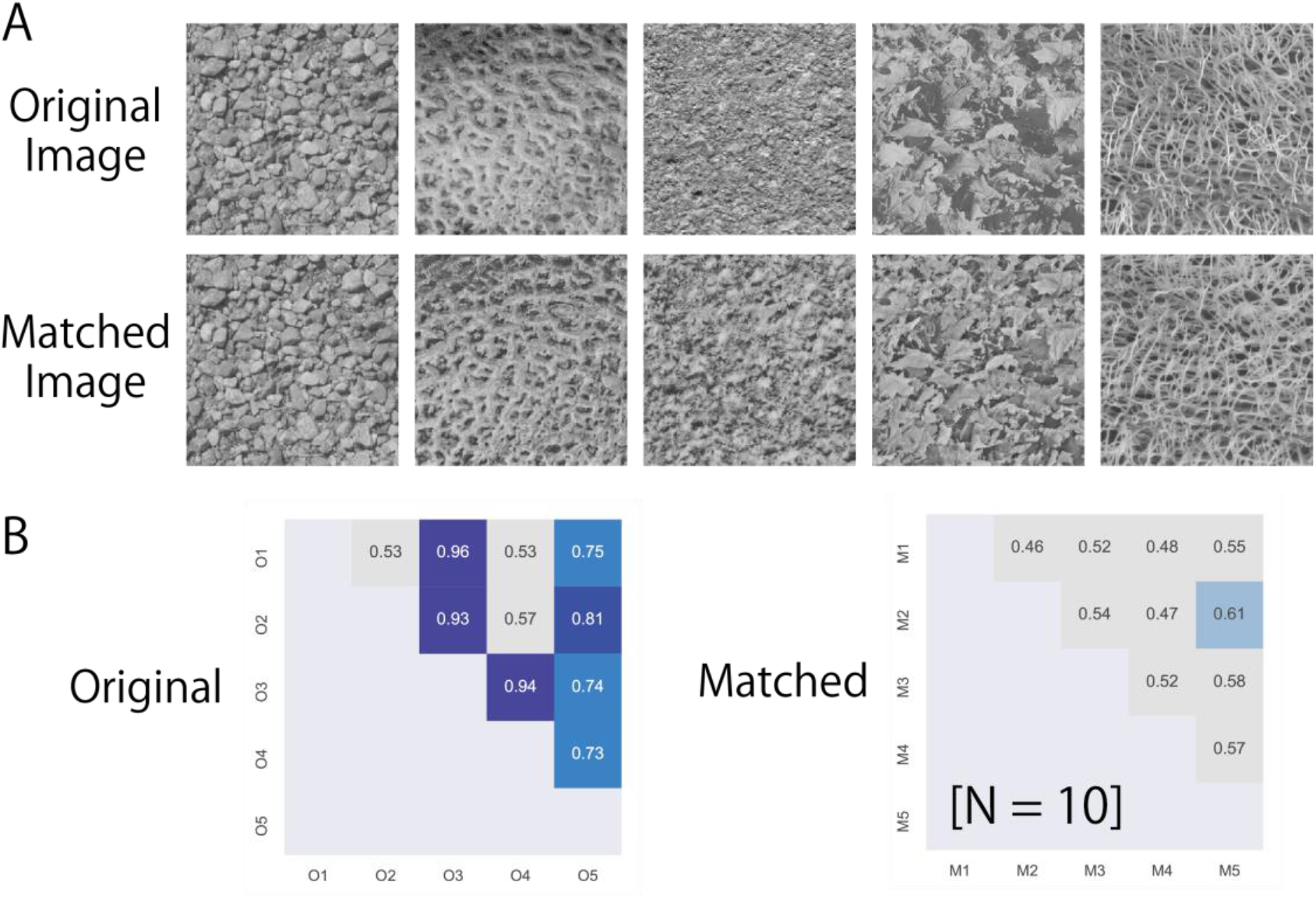
Original and matched NS stimuli. (A) Original NS images and histogram-matched NS images. Matched stimuli (M1-5) were based on different original images (NS/O1-5) but shared the subband histogram of NS1. (B) Results. Observers who could discriminate a few pairs of original NS stimuli could not discriminate NS-matched stimuli. The 95% confidence interval of the chance performance is 0.42-0.57 (uncorrected), and 0.38-0.62 (Bonferroni corrected).

It should be noted that this finding cannot be ascribed to the difference in spatial acuity between the two modalities. We made a gray-scale version of the histogram-matched images in the same size as the haptic stimulus (4 cm × 4 cm), and found them to be visually discriminable at the viewing distance of 3.4m where 1mm (haptic spatial resolution (Weinstein, 1968)) on the image subtended 1 minute of arc (maximum visual resolution of 20/20 observers).

### Texture discrimination experiment for 3D scanned surfaces

So far, the natural textures we used were made from visual images. To test the generality of our findings beyond this stimulus set, we checked whether subband matching could make metameric stimuli for haptic textures made from 3D scanned data of real textures. We selected eight surfaces of natural and artificial objects (Fig. 6) from a database of 3D scanned surfaces (Quixel Megascans). Note that our strategy here is to preserve the complexity and naturalness of the real world as much as possible, and thus these chosen objects have a highly diverse distribution in spatial scale and amplitude spectrum (e.g., the maximum carving depth Rz in Fig. 6). For example, chestnut stimuli are indeed much spikier than banana leaf stimuli (note that the image intensity of Fig. 6 is normalized on an image-by-image basis for the sake of pattern readability) and discrimination across newly printed natural haptic textures was easy by touch as well as by vision. Given that each texture has highly different frequency characteristics, exact matching of subband histograms across them (i.e., creating matched image by matching subband histogram of different textures as Fig. 5A) was hard or impossible. Thus, to test our hypothesis, this experiment separately examined difficulty in discrimination for each stimulus between the original texture and a histogram matched texture. The eight surfaces were 3D printed in the original scale and 1/3 scales. Then we created a histogram-matched version for each of the 16 stimuli by synthesizing the original one and a brown noise image, and tested whether human observers could discriminate each pair of the original and matched stimuli by passive scans. The results (group average shown in Fig. 6; see also Supplemental Table S2 for individual data) indicate that discrimination performance was not significantly different from chance (p>0.05) for the majority of the stimuli (11/16). These stimuli included not only image pairs visually very similar to each other (e.g., Concrete), but also those that look fairly different (e.g., Tile). Since we tested a wide range of textured surfaces, some of them contain large salient features (e.g., veins of banana leaf). For those textures, observers could use differences in these features in addition to those in surface textures to discriminate the original and matched stimuli, and this was probably the reason banana leaf and plaster showed above-chance performances. Overall, subband matching with natural haptic textures supports our conjecture that matching lower-order statistics make haptic textures metameric.

**Fig. 6.**
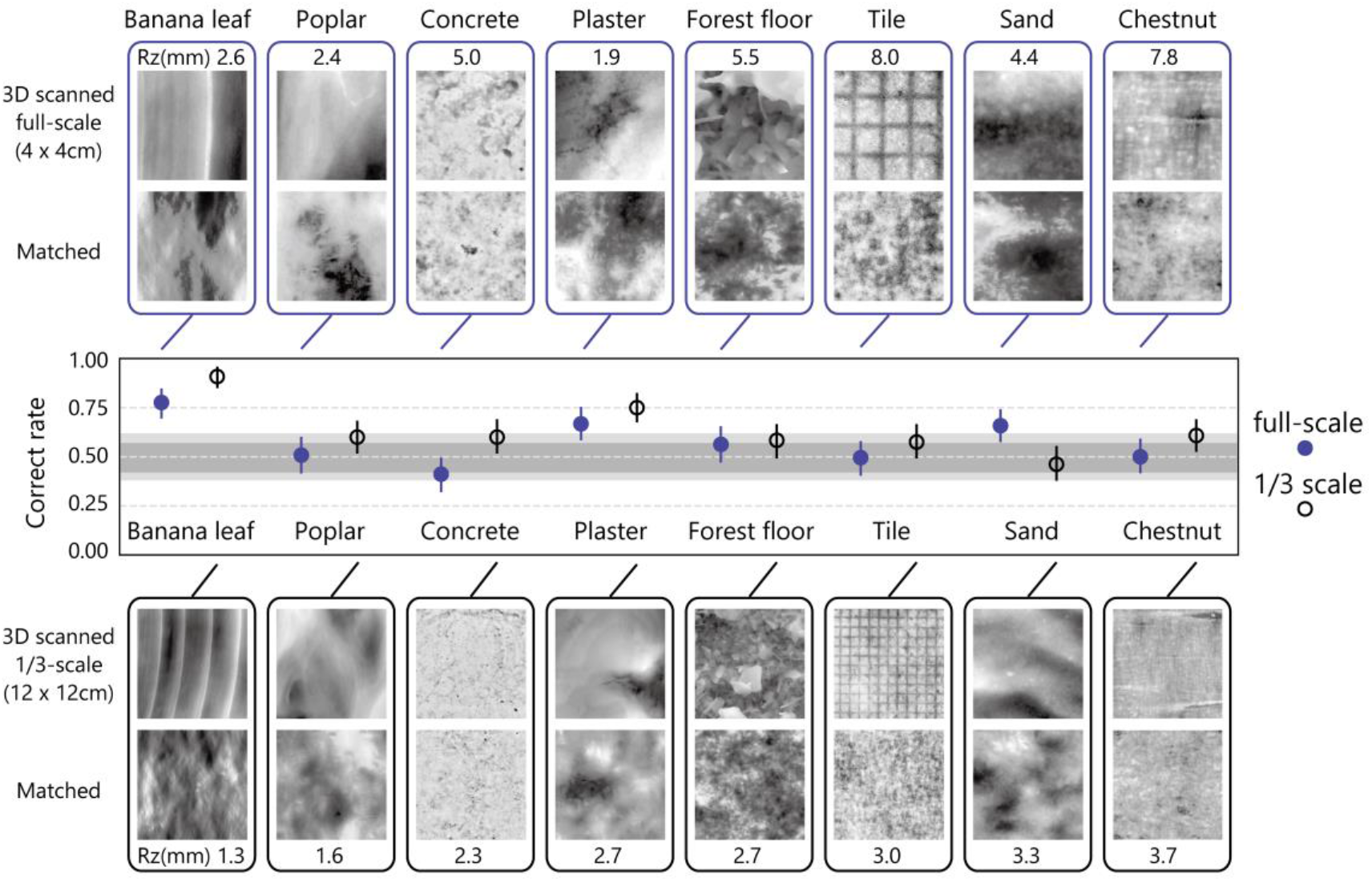
Original and matched textures made from 3D scanned surface. Images represent height maps of 3D scanned stimuli (top and third rows) and those of histogram-matched stimuli (second and fourth rows). Rz represents the maximum carving depth of 3D scanned surface. Darker means deeper, but the image intensity is normalized for the sake of pattern readability. Thus, a dark part of one image is not necessarily deeper than the brighter part of another image (e.g., the surface texture of banana leaf is much smoother and thinner than that of chestnut.) Each pair of 3D scanned stimulus and matched stimulus had the same subband histogram. In the graph, blue closed circles represent discrimination performance (proportion correct) in the ABX task with passive scans for each pair of full-scale stimuli (4×4 cm). Black open circles represent the performance for 1/3 scale stimuli (12×12 cm image scaled to 4×4 cm). Error bars denote a 95% confidence interval. Gray and light gray areas in the graph represent uncorrected and Bonferroni corrected 95% confidence intervals of the chance performance.

## General discussion

In this paper, our intent is to shed light on the similarities and differences in spatial-information processing mechanisms between vision and touch, the two major sensory modalities to perceive surface textures. The main question is whether these mechanisms are qualitatively similar in processing or whether they differ according to the accessible levels of stimulus features/statistics. We directly examined whether human observers can detect differences in higher-order texture statistics of 3D-printed stimuli, whose height patterns were manipulated with regard to image statistics. The key finding was that observers were unable to discriminate surface textures as long as the textures shared the same lower-order statistics (e.g., subband histograms). Note that these haptically metameric stimuli are visually discriminable, even we compensated for the difference in spatial resolution between the two modalities. Our tentative understanding here is that the tactile spatial pattern processor can take into account some statistics related to the local amplitude spectrum (level #1 and #2 in Fig. 1A), including the center frequency and bandwidth, but not those related to the phase spectrum or joint statistics (at level #3 in Fig. 1A). There are qualitative differences in spatial-information processing between vision and touch.

### Simulation of skin deformation and neural activity in periphery

We wonder whether the observed insensitivity to the higher-order statistics reflects processing characteristics of the central nervous system or simple information loss in the periphery, i.e., the elasticity of the skin and noisy sparse sampling by mechanoreceptors. To make our best guess of how the difference in spatial texture information is represented in peripheral neural activation, we simulated responses of tactile afferents using the computational model ‘TouchSim’ (Saal et al., 2017). The beauty of this model is that it can provide the responses of hundreds of distributed afferents on a millisecond scale, though it works under some simplified assumptions (e.g., it does not incorporate lateral sliding/forces). Using this model, we examined whether sufficient information for texture discrimination remains at peripheral stages.

We simulated the spike timings and spatial distributions of the afferents while the index finger pad scanned our subband-matched NS stimuli. We calculated the firing similarity across stimuli in terms of a spike distance metric (Victor and Purpura, 1997), following the original ‘TouchSim’ study (Saal et al., 2017). We found that the firing similarities between pairs of identical texture stimuli with independent neural noise were always higher than those between pairs comprising two different stimuli, regardless of the type of afferent (Fig. 7). This suggests that an ideal central encoder of the peripheral signals, which could fully utilize information including higher-order statistics, would be able to discriminate subband-matched stimulus pairs. Therefore, our finding of insensitivity to the higher-order statistics likely reflects processing characteristics of the central nervous system rather than information loss in the periphery mechanisms. It should be noted that unlike in the psychophysical ABX task, we did not rotate the texture when comparing the simulated neural responses between identical stimuli. This is because the firing similarity measure we used does not explicitly compute the similarity in higher-order statistics represented in terms of the relationship between neighboring afferent firings.

**Fig. 7.**
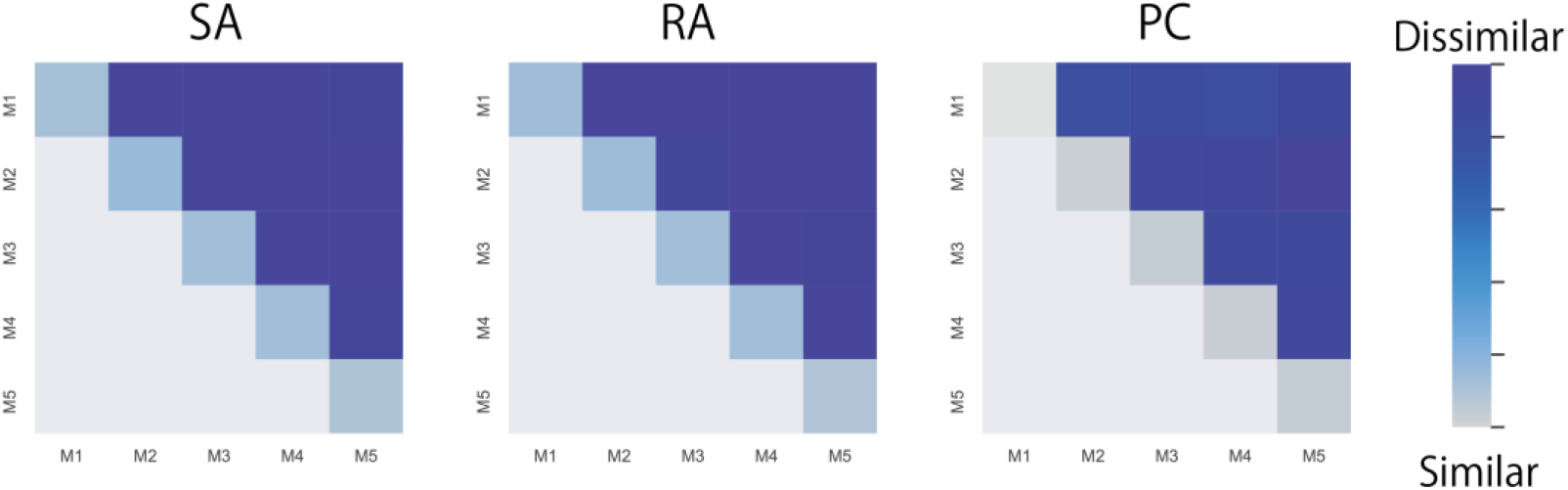
Similarity of neural firing patterns when the subband-matched stimuli were touched. The responses of tactile fibers of the index finger pad to NS-matched stimuli were simulated by the computational model ‘TouchSim’ (Saal et al., 2017). The similarity of simulated neural firing patterns was evaluated in terms of the Victor distance (Victor and Purpura, 1997). The distance between simulated firings was smaller (lighter colour) when the same stimulus (diagonal patches) was scanned than when two different stimuli were scanned (darker colour). See detail in method.

### Relationship with roughness perception

Roughness has been explored extensively and is recognized as a major feature in haptic texture perception (Bensmaia, 2009; Klatzky et al., 2013; Hollins et al., 1993, 2000; Tiest, 2010; Tiest and Kappers, 2006). Past studies have reported a variety of properties of roughness perception, including remarkable discrimination performance—we can detect even nanometre-scale differences (Skedung et al., 2013). Concerning the relationship with the current study, roughness perception can be ascribed to the tactile responses to lower-feature statistics, such as density or the deviation of surface elements (#1, #2 in Fig.1A). Industrial indexes of surface roughness, such as the arithmetical mean height of a surface (Ra) and maximum height of a surface (Rz) are also associated with lower-order statistics. As far as we are aware, the known characteristics of texture roughness perception do not conflict with our hypothesis that the tactile system is insensitive to higher-order feature statistics. In addition, not only us but other groups as well noticed this haptic metamerism of the subband matched stimuli. For instance, a group of tribology researchers 3D printed surface textures with random topography, and their stimuli were very similar to ours. Their recent conference paper reported a low discrimination ability in human observers (Sahli et al., 2019), which is also consistent with our results and theory.

Though our observers could use texture differences in dimensions other than roughness to accomplish the ABX discrimination task with our complex stimuli, we do not exclude a possibility that they mainly relied on what previous studies called roughness in performing the discrimination task. We therefore considered whether the current results can be accounted for by previously proposed neural indexes of roughness perception. We computed two neural roughness indexes—the mean impulse rate of high-frequency PC afferents (Bensmaia & Hollins, 2005; Hollins & Bensmaia, 2007) and spatial variation of low-frequency SA afferents (Connor et al., 1990, 1992)—from the output of ‘TouchSim’. Note that these indexes do not take into account higher-order statistics, so we did rotate the texture and add spatial jitter derived from contact position variability when repeating the simulated neural responses. We found the simulated indexes to indicate similar or even higher discrimination performance for NS-original and NS-matched stimuli compared to the observed behavioural performance (Fig. S4). The predictions of these indexes might be improved by adjustment of noise parameters. Further investigation is warranted to determine how well these or other neural roughness indexes can explain tactile perception of complex texture patterns like the ones we introduced.

There are textures that look identical (visually metameric) but can be discriminated by touch due to differing microstructures that are too small for the eye to resolve. However, this is not inconsistent with our conjecture, since such textures with microscale roughness do not have specific higher-order spatial structures and are presumably detected through temporal vibration patterns of skin responses (Bensmaia & Hollins, 2005; Hollins & Risner, 2000; Johnson et al., 2002).

### Relationship with shape perception

Although our findings indicate that tactile *texture* perception is insensitive to higher-order statistics, they do not exclude a possibility that tactile *shape* perception has sensitivity to some higher-order features. Tactile shape perception shows high sensitivity to stimulus orientation. Edge orientation acuity was reported to be around 20 degrees (Bensmaia et al., 2008; Olczak et al., 2018) or even smaller (Pruszynski et al., 2018). Furthermore, human touch can discriminate more complex spatial patterns such as letters of the alphabet. One may interpret this result to suggest the ability of the tactile shape perception to discriminate some spatial phase differences. Indeed, neurons in area 2 (Fitzgerald et al., 2006; Yau et al., 2013) and parietal opercular cortex (Yau et al., 2009) show sensitivity to higher-order shape features (i.e., particular curvatures). However, tactile shape perception and tactile texture perception have been studied nearly independently by using different types of stimuli. Shape perception has been tested with simple/local stimuli such as raised simple line/curvature patterns, while texture perception has been tested with complex/global stimuli, including natural textures. While shape perception focuses on a specific location in the stimulus, texture perception should grasp the holistic statistical nature of the field under crowding conditions as in the current experiments. We therefore do not assume that texture perception can be explained by the same mechanism as that for shape perception, and this difference may be reflected in the above-chance discrimination performance of a few matched natural haptic texture pairs. Indeed, shape processing and texture processing may be segregated in somatosensory cortex (e.g., Lesions in parietal opercular cortex are known to impair shape recognition but not roughness recognition (Roland, 1987)). At present, we have no clear evidence of the contribution of a neural mechanism sensitive to higher-order features (curvature) to surface texture perception (see also Fig S5).

### Relationship with braille reading

Experienced human observers often show a remarkable ability to read braille characters by touch, and this tactile character recognition is very similar in performance to visual character recognition when the image resolution is matched between the modalities (Apkarian-Stielau & Loomis, 1975; Craig, 1979; Loomis, 1981, 1982; Phillips et. al., 1983). Our conjecture that tactile texture perception is unable to use higher-order spatial statistics may appear incompatible with such tactile character reading. However, we do not believe it is. First of all, many characters differ from one another in terms of amplitude spectra. In the case of braille, among 325 pairs of the 26 letters of the English alphabet, only four pairs have identical dipole (second-order) statistics and thus identical amplitude spectra (Julesz, 1973). All of them are vertically flipped pairs. Within each pair, the two characters are sometimes confused with each other, but not very often (Loomis, 1982). Successful discrimination of isodipole braille characters however should be regarded as tactile form perception based on position and/or orientation of specific features. In contrast, what we examined here is tactile texture perception based on the image statistics sampled at multiple locations within the texture. Note also that our participants attempted to identify two textures in different overall orientations. Under this condition, discrimination of isodipole braille characters would be logically impossible. Therefore, the ability to read tactile characters does not counter our argument about the computational limitation in tactile texture processing.

### Future directions

To analyze the sensitivity of the tactile system to image statistics using a method similar to that used in vision research, we transcribed gray-scale visual images into surface depth maps. Due to the difference in the sensing process, however, we can only assume a rough correspondence of the sensor response pattern between the two modalities. To overcome this limitation, we used ‘TouchSim’ (Saal et al., 2017) to evaluate our hypothesis in this study, but more direct evaluation would be necessary in future.

Overall reversal of the signal sign (contrast polarity) affects some lower-order statistics (e.g., the pixel intensity histogram (first/lower-order statistics)), but not the local amplitude spectrum. While the visual system responds to positive (white) and negative (black) elements in an asymmetric way (e.g., Chubb et al., 2007), the tactile system may do so much more strongly. A sign reversal swaps convex elements with concave ones, thereby introducing a large change in skin deformation patterns (e.g., pin or hole elements). Indeed, we found that human observers could discriminate the original and the sign-reversal version for our NS stimuli (Fig. S6). Note that this result does not conflict with our general conclusion of haptic insensitivity to higher-order statistics, since negative-positive inversion stimuli share amplitude spectrum but differ in other lower-order statistics. One way to cope with this kind of situation with the local amplitude spectrum is to consider the low-pass characteristics of elastic skin. By inserting a non-linear mapping process, precipitous height patterns could be more realistically transferred to sensor response patterns. This procedure indeed produces a significant difference in the amplitude spectrum of the sensor responses between the original and sign-reversed version.

### Concluding remarks

Recent 3D printing technology allows us to control the spatial pattern of tactile stimuli as accurately and flexibly as in the case of visual stimuli. This powerful methodology makes it possible to apply a variety of experimental paradigms developed in vision research to tactile research. As we demonstrated here, 3D printers will be powerful tools for future investigations of tactile spatial computation.

Several studies have investigated the relationship of touch with vision, as this study did. While several past influential studies (Amedi et al., 2001; Kitada et al., 2006; Merabet, 2004; Yau et al., 2009; Zangaladze et al., 1999) emphasized the similarities between the two modalities, our study rather highlighted the differences (see also Lederman et al., 1990; Whitaker et al., 2008). By doing so, we could clarify in what way spatial sensation by touch is different from spatial sensation by vision. While many people may share the intuition that tactile texture sensation is qualitatively different from visual texture sensation, here we showed, for the first time, that the qualitative differences arise from (in)sensitivity to higher-order image statistics.

## Methods

### Generating visual texture stimuli

Since our tactile stimuli were created by 3D-printing according to a height map, we dealt with the height map as a visual image and controlled its image statistics as in the visual texture literature. The image we used was either artificial Gaussian band-pass noise or a natural scene texture of 256×256 pixels. For the artificial noise, we first applied the two-dimensional fast Fourier transform to a white noise image and extracted specific spatial frequency components by using a two-dimensional Gaussian band-path filter. There were two stimulus conditions of filter parameter manipulation: the center frequency (CF) and bandwidth (BW) conditions (Fig. 2A, top and center). The CF condition used five center frequencies of the band-path filter (2, 4, 8, 16, and 32 cycles/image) while keeping the BW 1.5 octaves. The BW condition used five filter bandwidths (0.5, 1.0, 1.5, 2.0, and 2.5) while keeping the CF 8 cycles/image. In addition to the artificial noise, we used the natural scenes (NSs). For this NS condition, five images were chosen from the natural texture category of the McGill Calibrated Colour Image Database (Olmos and Kingdom, 2004). The intensity level (the mean and standard deviation) were normalized and equalized across images (Fig. 2A, bottom). For 3D scanned stimuli condition, eight height maps (Banana leaf, Poplar bark, Concrete surface, Plaster surface, Forest floor, Tile surface, Sand floor, Chestnut bark) were chosen from Quixel megascans library. Each height map (i.e., exr image) had 8M pixels for 2m×2m (or 1m×2m) scanned surface, and the contrast difference between complete black (0) and complete white (1) in an image describes a height (thickness of stimuli; black means deep) difference of 20 cm. Each image was trimmed to 4cm×4cm (3D printed in original scale) and 12cm×12cm (3D printed in 4cm×4cm as 1/3-scale). Note that images in Fig. 6A were displayed with automatic contrast control for the sake of readability and original scale was not kept. Indeed, height map image of Banana leaf had much lower contrast compared to that of Chestnut bark.

### Generating tactile texture stimuli

Tactile stimuli were custom-built for the experiment by using a 3D printer (16-micrometer resolution) (Objet 260 Connex3, Stratasys, USA) with transparent plastic-like material (VeroClear-RGD810, Objet, USA). Each visual texture stimulus was converted into a 3D model by taking intensity values as a height map. The printed object was 40×40×10-12 mm. The contrast difference between complete black (0) and complete white (255) in an image was transcribed to a height (thickness of stimuli; black means deep) difference of 2 mm. Printing accuracy, measured with a wide-area 3D measurement system (VR-3100, KEYENCE, Japan), was within 0.056 mm on average. For full-scale 3D scanned stimuli, the object was printed in original scale, i.e., 40×40mm surface with different maximum carving depth (Banana leaf: 2.6, Poplar bark: 2.4, Concrete surface: 5.0, Plaster surface: 1.9, Forest floor: 5.5, Tile surface: 8.0, Sand floor: 4.4, Chestnut bark: 7.8mm). For 1/3-scale 3D scanned stimuli, the object was printed in 1/3 scale in three dimensions, i.e., 40×40mm surface with different maximum carving depth (Banana leaf: 1.3, Poplar bark: 1.6, Concrete surface: 2.3, Plaster surface: 2.7, Forest floor: 2.7, Tile surface: 3.0, Sand floor: 3.3, Chestnut bark: 3.7mm). Prior to the experiment, the surface of the stimuli was lightly covered with baby powder (Baby Powder, Johnson & Johnson, USA) to ensure constant contact between the finger and stimuli by avoiding large stick-slips.

### Generating tactile vibration stimuli

A piezoelectric actuator (MU120, MESS-TEK, Japan) was used as stimulator to reproduce the line scan of the texture height profile swept along a horizontal line. The stimulator normally deformed the skin with a maximum of 800 N force vibrated by a position control method so that it could accurately produce the required displacement with a tolerance of few nanometers. The diameter of the stimulator was 12.0 mm, and its edges were separated from the rigid surround of the metal boards by a 1.0-mm gap (following (Verrillo, 1963)). The rigid surround limits the spread of surface waves of the skin. The stimulator always contacted the finger throughout the experiment. The line on the textured surface to be converted into vibration was randomly chosen for each trial. Note that due to actuator limitations, the original height difference was linearly reduced to fit within 120 μm (roughly 1/10 scale).

### Participants

Thirty-six naïve observers and two of the authors (13 males) with normal tactile sensitivity (by self-reports), aged from 21 to 46 years (32.2±7.73) participated the experiments. All gave informed consent approved by the NTT Communication Science Laboratory Research Ethics Committee, and all procedures were conducted in accordance with the Declaration of Helsinki.

### Procedures

#### ABX experiment with CF, BW, and NS modulation

Groups of ten observers participated for each passive, active, static, and vibration condition, with partial overlaps of observers across conditions. An observer sat at a table and placed the index finger of the right hand on home position with their right arm on an arm rest. They performed experiments with eyes open to maintain their arousal level, but they could not see the tactile stimuli, the equipment, nor experimenter, which were occluded by a black curtain.

In general, observers touched three stimuli, A, B, and X, each for one second, where X was a rotated version of A or B, and they verbally reported which one was X, the first or second. Since X was rotated, observers could not perform feature matching using trivial keys. Paired stimuli (A and B) were randomly chosen from five stimuli of the same modulation (CF, BW, or NS). This procedure is called an ABX task (Kingdom & Prins, 2010), and one of the advantages of this task is that it is less affected by labeling problems. If the observers are asked to directly evaluate the similarity between paired stimuli, these stimuli must be labeled, and their responses may be influenced by labeling difficulties and/or confusion between labels. On the other hand, with the ABX task, the observer can report the relative similarity between X and A and X and B, even if A and B are not clearly labeled. There were four different touching mode conditions. Other than in the vibration condition, the experimenter set three predetermined stimuli (A, B, X) on a linear stage (ERL2, CKD, Japan) before each trial started. By automatically moving the stage, the experimenter was able to guide the three stimuli directly beneath the finger. Thus, observers did not have to move the wrist to touch them. In the passive scan condition, each trial started with the experimenter’s ‘ready’ call. An observer put their right index finger at the rest position at the right edge of the linear stage and pressed the start button on the PC monitor with their left hand to trigger the stage movement. The linear stage started to move under observers’ right index finger from left to right for approximately 1.5 seconds with a speed of 40 mm/s so that the rightmost one of three stimuli swiped the finger for one second. After a one second pause, the stage automatically moved again for the second stimulus to swipe, stopped, then moved again for the third stimulus to swipe. After the third scan, observers lifted their fingers and made a binary verbal report as to which of the first two stimuli (the first or the second) was the same to the last one. The stage moved back to its initial position and the experimenter changed the stimuli on the stage for the next trial. In active scan condition, observers lifted their right index finger above the first stimulus position and pressed the start button. They freely scanned the stimulus surface for one second and lifted their finger again. Then, the stage started to move and the second stimuli came right under their finger. Observers never touched the stimuli when the stage was moving; they touched them three times each for one second when the stage had stopped. In the static touch condition, observers put their fingers on static (not moving) stimuli three times each for one second, in similar time course to other conditions. In the vibration condition, the stage and stimuli were replaced with the piezoelectric actuator. Observers placed their right index finger on the actuator and pressed the start button on the monitor with their left hand. After about one second, the actuator vibrated three times each for one second with 1.5-second gap. Only in this condition did observers report by button clicking instead of verbally. To mask any subtle sound made by the actuator, observers wore ear plugs and white noise was played continuously from headphones throughout this condition.

There were three kinds of modulation (CF, BW, and NS), five modulation gradations, four touching modes (passive scan, active scan, static touch, and vibration), and 12 repetitions for each combination. In total, 360 trials were conducted for each touching mode, and sessions were roughly divided into ten blocks. No block lasted longer than 15 minutes, with at least a 10 minutes break between successive blocks. Within each block, the kind of modulation and the gradations were randomized and the touching mode was fixed. Each observer completed the main session in 90-120 minutes, and addition of practice sessions and breaks doubled the total duty time.

#### ABX experiment with NS original and NS matched modulation

Ten observers participated. The equipment and procedure were almost identical to those of the passive scan condition in the ABX experiment, except for the stimulus. The experiment was conducted with two kinds of modulation (original and matched NS), five modulation gradations, one touching mode (passive scan), and 12 repetitions for each combination. Each observer completed the session in 70-90 minutes (except breaks).

#### ABX experiment with 3D scanned stimuli and matched stimuli

Ten observers participated. The equipment and procedure were almost identical to those in the passive scan condition in the ABX experiment, except for the stimulus pair (i.e., how A and B were chosen). In other experiments, A and B were chosen from the same kind of modulation (e.g., A = NS1, B = NS2). In this experiment, A and B were chosen from the same scale of same object (e.g., A = 3D scanned concrete surface, B = Matched concrete surface). The experiment was conducted with two kinds of origin (3D scanned stimuli and matched stimuli), eight different object, two different scale (full-scale and 1/3 scale), one touching mode (passive scan), and 12 repetitions for each combination. Each observer completed the session in 70-90 minutes (except breaks).

### Logistic regression analysis based on amplitude differences

To investigate whether the discrimination performance can be explained by the amplitude differences in the textures, we used a logistic regression analysis. As shown in Fig. 4A, the amplitude of each spatial component was calculated on a log scale and sampled to ten points in 0.7-octave intervals. The difference between the amplitude spectra of paired textures was calculated at each sampled point. We used the ten values as predictor variables for each discrimination performance as follows.

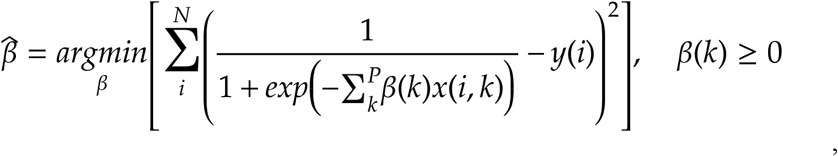

where x is the predictor variable, β its coefficients, y the human discrimination accuracy for each pair, N the number of pairs, and P the number of the predictor variables. We used Matlab function *fmincon* for the optimization. By the sparse logistic regression, five of the ten predictor variables survived, and were used in estimating the performance shown in Fig. 4B.

### Generating the subband-matched images

To test if textures with identical lower-order (subband) statistics were haptically indistinguishable, we made five images with identical subband histograms (Fig. 5A, bottom panels M1-M5) by using a texture synthesis algorithm (Heeger & Bergen, 1995). In this algorithm, the steerable pyramid transform is utilized. Specifically, each image was decomposed into several spatial-frequency and orientation bands by convolving the image with spatially oriented linear filters and by subsampling. Using the transformed subbands, this algorithm synthesizes the texture by matching the histograms of transform coefficients of the seed image to those of the target image. We used original image 1 (NS/O1) as the target for the synthesis. The original algorithm uses a white noise image as the seed image. However, since we aimed to make a set of textures while preserving the higher order statistics of each original texture (O1 – O5), we used the original texture as the seed image. To make the matched images in Fig. 5, we adopted four spatial frequencies bands and four orientation bands.

The texture synthesis algorithm of Heeger & Bergen (1990) was recruited to also make the matched height maps of 3D scanned stimuli (Fig. 6A). We used each original 3D scanned height map as the target for the synthesis. Since the subband distribution of the tactile textures was spatially biased than the visual textures in the previous experiments, we adopted more bands for the synthesis, specifically six spatial frequency bands and six orientation bands, and used a brown noise image as the seed image to decrease the difference of subband histograms between the original and synthesized images.

### Simulation

We simulated spatio-temporally distributed firing patterns of three different types of tactile afferent by using the computational model “TouchSim”, which can reproduce major response characteristics that have been clarified by previous research (Saal et al., 2017). The original parameters of the model were based on measured spiking data obtained with monkeys. The simulated responses of afferents closely match the known spiking responses of actual afferents (both precise millisecond spike-timings and firing rates) to various classes of stimuli (for example, vibrations, edges, and textured surfaces).

Responses to NS matched stimuli were simulated. The stimulus was scanned across the skin. The contact area was defined as a rectangle (20-mm length and 10-mm width) and the resolution (input spacing, defined as pin spacing in ‘TouchSim’) was set to 0.1 mm. The skin contact area was indented at the center of the index fingertip with the averaged depth of 1 mm, moved across the stimuli at a speed of 40 mm/s for one second. Realistically distributed afferents (288 SA, 569 RA, and 102 PC in index fingertip ‘D2d’) were simulated with a 1-ms resolution. The simulation was repeated 12 times for each stimulus, with afferent distribution, stimulus contact area, and scan direction fixed.

Since we simulated temporally and spatially distributed firings, higher-order statistical information, if any, should be embedded in the firing pattern. To quantitatively test whether this pattern is similar when touching the same stimulus compared to when touching different stimuli, we conducted metric space analysis. In particular, we calculated the Victor distance (50) following the previous ‘TouchSim’ study (Saal et al., 2017). This analysis enables us to compare the similarity of the timing of the spikes by introducing cost parameter q, which we set to 100 (corresponding to 10 ms) in this study. For each afferent model, the Victor distance was calculated between all pairwise combinations of the simulated trials for five stimuli. Obtained distances were normalized by the total number of spikes in the pair and then averaged for the same kind of afferent model and for the same pair of stimuli (Fig. 6).

## Author contributions

SK, MS, Conception and design, Acquisition of data, Analysis and interpretation of data, Drafting and revising the article; SN, Conception and design, Drafting and revising the article

## Ethics

Human observers: Forty healthy people participated these experiments. All observers provided written informed consent in accordance with the Declaration of Helsinki. The ethics committee at NTT Communication Science Laboratory Research approved the study.

