## Supplemental File for "Haptic metameric textures"

### Supplementary material

*MDS Experiment with CF, BW, NS modulation.*

Ten observers participated, who partially overlapped the group of the ABX experiment. The equipment and procedure were almost identical to those in the passive scan condition in the ABX experiment, except that observers touched only two stimuli and rated their perceived dissimilarity between 0 (identical) to 100 (very different). Ratings were provided by mouse clicking on a ruler on the display. The pairs of stimuli were randomly chosen from 15 (five each from the CF, BW, and NS stimuli, and each stimulus was presented once in the first and the second intervals for every pair. The dissimilarity data consisted of two ratings per pair per observer. Before the experiment, observers practiced ratings for ten trials, and these data were not included in the analysis. Each observer completed the main session in 90-120 minutes, and addition of practice sessions and breaks doubled the total duty time.

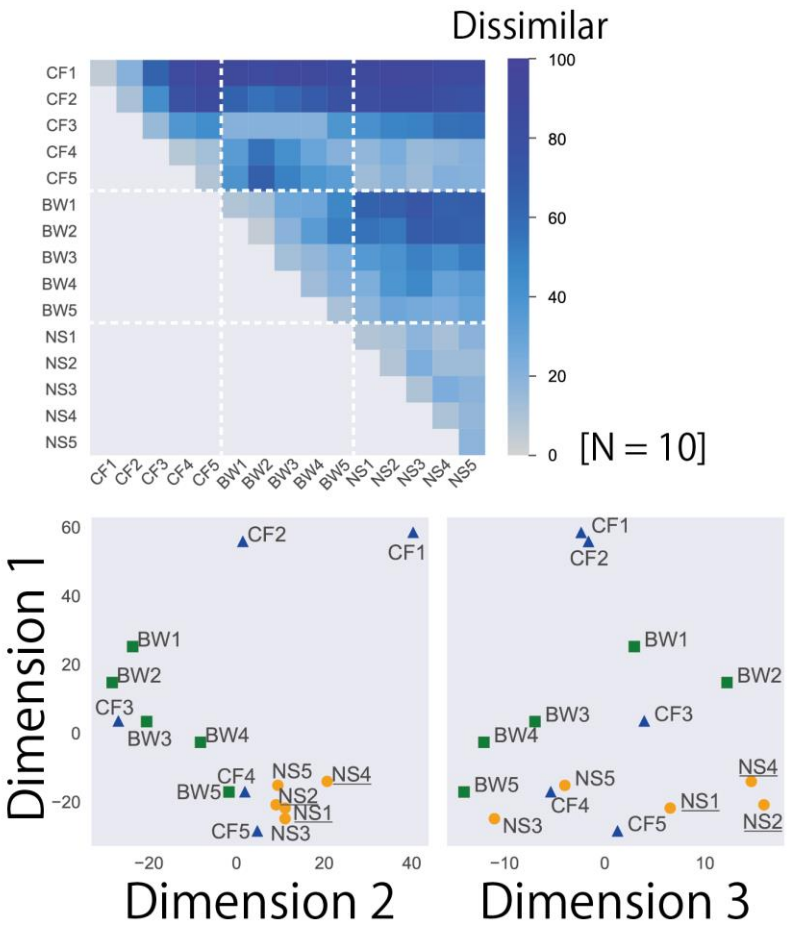


Fig. S1. Results of MDS experiment. Three-dimensional tactile space based on perceived similarities among CF, BW, and NS stimuli; the closer the points in the map are to each other, the more similar the surfaces are perceived. Underlined stimuli were indistinguishable ones.

*ABX experiment with four different touching modes*

To consider the effect of the touching mode, we used four different ones. Ten observers participated for each mode, who were partially overlapped across modes. The full set of results is shown in Fig. S2. The results for NS stimuli are the same as those reported in the main manuscript.

The differences between the passive and active scan modes for CF and BF stimuli were roughly similar to that for NS stimuli. Performance with the active scan mode (0.96 for CF; 0.85 for BW on average) was similar or slightly better than that with the passive scan (0.91 for CF; 0.76 for BW). In the static touch condition, temporal information was limited. In the vibration condition, spatial information was limited. Since the tactile system shows drastic spatial and temporal summation characteristics with subthreshold stimuli (Gescheider et al. 1999, 2005), one might expect that these limits would improve the texture discrimination performance. The observed performance however was even worse than in the scan conditions for all stimulus sets, suggesting the importance of both spatial information and temporal information in haptic texture discrimination.

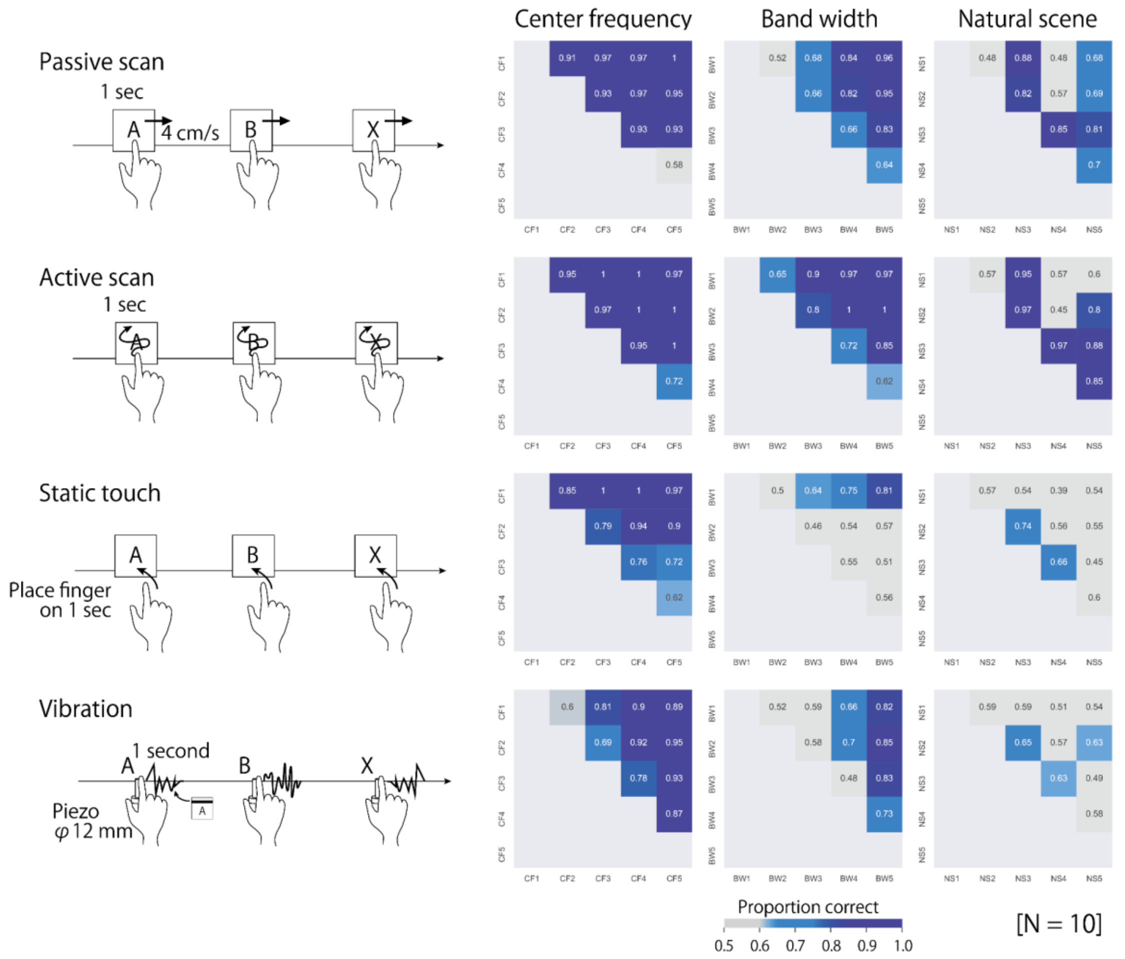


Fig. S2. All combinations of modulations and touching modes. Note that different touching modes were not necessarily performed by the same observers.

*MDS experiment with NS original and NS matched modulations.*

Ten observers participated. The equipment and procedure were almost identical to those of the passive scan condition in the MDS experiment, except for the stimulus. The experiment was conducted with ten stimuli (five original and five matched NS stimuli), one touching mode (passive scan), and two repetitions for each combination. Each observer completed the session in about 60 minutes (except breaks).

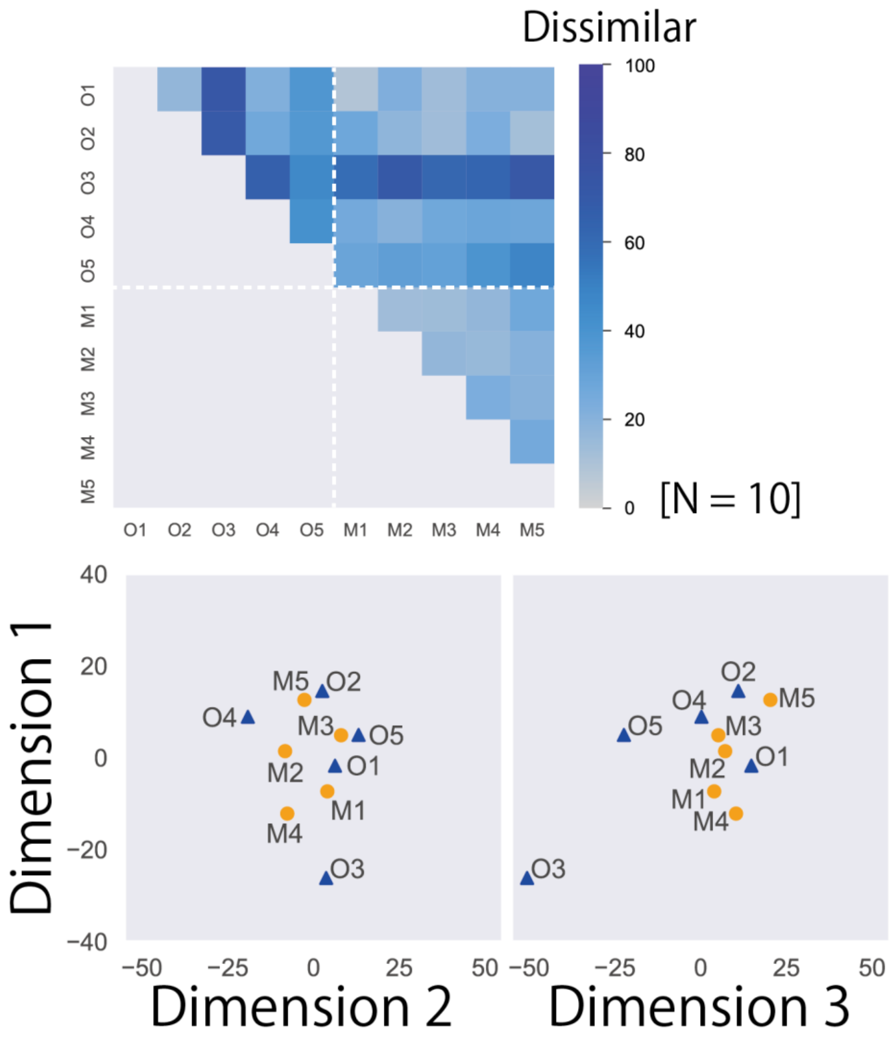


Fig S3. Three-dimensional tactile space based on perceived similarities among original and matched NS stimuli. Matched stimuli (M1-5) have the same subband histogram as original NS1 (here named O1), which is well represented by their being distributed around O1.

*Simulating neural indexes based on 'TouchSim'*

Since roughness is a major feature in haptic texture perception, we calculated two potential neural codes of roughness perception using the simulated firing data. One is local spatial variation in the firing rate in the SA channel [Connor, Hsiao, Phillips, & Johnson, 1990]; the other is intensity (mean firing rate) in the PC channel [Hollins & Bensmaia, 2007]. To test how well these codes can account for our results, the accuracy of an ideal observer to perform our ABX task using these neural codes was calculated and compared with behavioural accuracy. The local spatial variation of each SA afferent was calculated as the mean absolute differences in the firing rate in comparison with neighboring afferents within a 3-mm distance. For a trial of each stimulus condition, we calculated this variation for each SA afferent and then averaged across all the afferents. The PC's mean firing rate was measured by summing the total firings. For a trial of each stimulus condition, we calculated this rate for the response of each PC model afferent and then averaged across all the afferent. To estimate the predicted accuracy of the ABX task, we computed the probability of each index being larger when scanning the same stimulus repeatedly than when scanning different stimuli. We repeated the whole processes 12 times to mimic our experimental condition. The calculated indexes varied across repetitions due to the model's internal noise. Though these indexes do not take higher-order statistics into account, the simulated indexes suggested similar or even higher discrimination ability of NS stimuli compared with behavioural performance (Fig. S4). Since the absolute level of estimated performance could vary depending on the decoding process (here we simply compared raw index values without assuming additional noise or filtering before comparison), what is important is similarity in response pattern, rather than in absolute value, with human accuracy. In this respect, the result for the local spatial variation of the SA channel (square symbols) is relatively closer to the behavioural one (Pearson's r=0.82) than that for the mean firing rate of the PC channel (circle symbols) is (r=0.28).

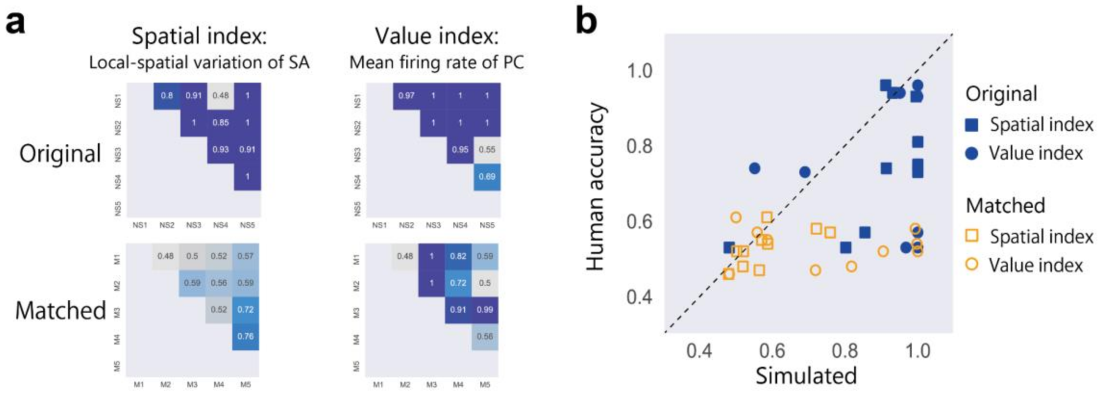


Fig. S4. Simulated performance of ABX task based on computational model 'TouchSim' [Saal et al., 2017]. (**a**) Estimated performance of ABX task with conventionally proposed neural indexes. (**b**) Comparison between observed performance (vertical axis, data from Fig. 5b) and estimated performance (horizontal axis, data from panel **a**).

*Individual differences in ABX and MDS experiments*

Some observers distinguished some pairs of matched NS stimuli, though the trend was not common across observers.

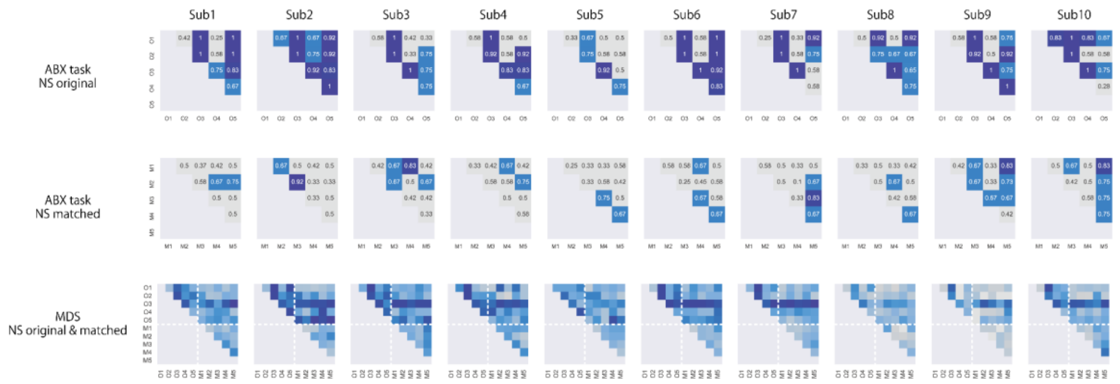


Fig. S5. Individual results of ABX task and MDS with original and matched NS stimuli.

*ABX experiment with NS and NS inverse modulation.*

Switching black and white in an image results in the same amplitude spectrum in image analysis, while it switches convex and concave (changing pins to holes) in 3D printing. This manipulation would induce huge differences in skin deformation. Though our results suggested haptic discrimination performance is correlated with differences in the amplitude spectrum, this black and white manipulation may result in exceptions. To test this, we printed NS inverse stimuli using black and white reversal images of NS images and measured the discrimination performance for NS stimuli and NS inverse stimuli. The equipment and procedure were almost identical to those in the passive scan condition in the ABX experiment, except for the stimulus pair (i.e., how A and B were chosen). In other experiments, A and B were chosen from the same kind of modulation (e.g., A = NS1, B = NS2). In this experiment, A and B were chosen from the same modulation gradation with different kinds of modulation (e.g., A=NS1-original, B=NS1-inverse). Ten observers participated. 12 repetitions for each combination. Each observer completed the experiment in about 30 minutes (except breaks). The results showed that indeed human observers could discriminate the original and inverted patterns of our NS stimuli. This result however does not necessarily imply haptic sensitivity to higher-order statistics, since a polarity reversal changes the pixel intensity/depth histogram (first-order statistics).

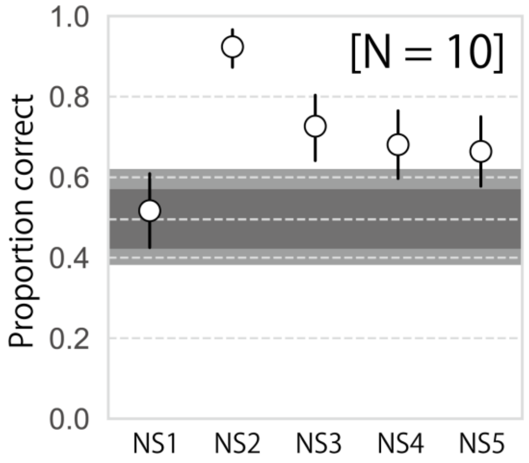


Fig S6. Performance of ABX task with original and inverted NS stimuli. NS2-original could be discriminated from NS2-inverse above chance level, even though these two stimuli share the same amplitude spectra. Error bars denote 95% confidence interval. Gray area and light gray area in the graph represent uncorrected and Bonferroni corrected 95% confidence interval of the chance performance.

*Individual performance of each observer*

We tried to statistically check whether individual observers' response in discriminating matched stimuli had any possibility to exceed chance level performance. To do so, we calculated 95% (5% threshold) and 99.5% (corrected for multiple comparison) confidence interval of the chance performance (0.5 in ABX task) by binomial distribution. Note that we averaged the performance of all paired conditions (all 10 cells in one graph in Fig. 5B) for each observer, since the repetition number is low. Thus the distribution of 120 trials (12 repetitions of 10 pairs) was calculated. We found that for Matched condition, only 1 observer (out of 10) showed above 95% (CI=0.42-0.57) and 99.5% (0.38-0.62) range performance, while most of them was not. In original NS condition, this was not the case, and all 10 observers overshot the CI range.

Table S1: Analysis of results with original and matched NS stimuli. Ratio of the number of observer who could discriminate stimuli better than 95% or 99.5% range of chance performance.

| | 5% Threshold | 5% Threshold (w/ Bonferroni correction) |
|---|---|---|
| Original NS | 10 / 10 | 10 / 10 |
| Matched NS | 1 / 10 | 1 / 10 |

We also conducted this analysis on experiment with natural tactile textures. When averaging the performance of all stimuli, a significant number of observers showed above CI performance. Still, we are reluctant to say that observers can robustly discriminate these (scanned and matched) stimuli, since the ratio dropped when we exclude samples that contain large-scale localized features (e.g., vein that resembles a shape rather than a texture). In addition, there is no positive evidence of better discrimination with full-scale stimuli than 1/3 scale stimuli, which can be expected from the possible benefit of the "real" scale if there is any.

Table S2. Analysis of results with natural tactile texture. Ratio of the number of observers who could discriminate stimuli better than 95% or 99.3-4% (corrected for multiple comparison) range of chance performance.

| | 5% Threshold | 5% Threshold (w/ Bonferroni correction) |
|---|---|---|
| full-scale (All) | 4 / 10 | 2 / 10 |
| 1/3 scale (All) | 9 / 10 | 5 / 10 |
| full-scale (w/o Banana) | 2 / 10 | 2 / 10 |
| 1/3 scale (w/o Banana) | 5 / 10 | 3 / 10 |
| full-scale (w/o Banana Plaster ) | 2 / 10 | 1 / 10 |
| 1/3 scale (w/o Banana Plaster ) | 2 / 10 | 0 / 10 |
